# Mobile stamens enhance pollen dispersal by scaring floral visitors away

**DOI:** 10.1101/2022.07.06.498951

**Authors:** Deng-Fei Li, Wen-Long Han, Susanne S. Renner, Shuang-Quan Huang

## Abstract

Animal-pollinated plants have to get pollen to a conspecific stigma while protecting it from getting eaten. We provide experimental evidence that touch-sensitive stamens function in (i) enhancing pollen export and (ii) reducing pollen loss to thieves. Stamens of *Berberis* and *Mahonia* are inserted between paired nectar glands and when touched by an insect’s tongue rapidly snap forward so that their valvate anthers press pollen on the insect’s tongue or face. We immobilized the stamens in otherwise unmodified flowers and studied pollen transfer in the field and under enclosed conditions. On flowers with immobilized stamens, the commonest bee visitor stayed up to 3.6x longer, yet removed 1.3x fewer pollen grains and deposited 2.1x fewer grains on stigmas per visit. Self-pollen from a single stamen hitting the stigma amounted to 6% of the grains received from single bee visits. Bees discarded pollen passively placed on their bodies, likely because of its berberine content; nectar has no berberine. Syrphid flies fed on both nectar and pollen, taking more when stamens were immobilized. Pollen-tracking experiments in two species showed that mobile-stamen-flowers donate pollen to many more recipients. These results demonstrate another mechanism by which plants simultaneously meter out their pollen and reduce pollen theft.

**Highlights:**

- Stamens that snap forward when triggered by a flower visitor may serve to meter out pollen, scare away pollen thieves, or place pollen more accurately.
- We tested these hypotheses by experimentally immobilizing all six stamens in numerous flowers of *Berberis* and *Mahoni*a species in the field and under enclosed conditions.
- In flowers with immobilized stamens, the commonest bee species stayed up to 3.6x longer, yet removed 1.3x fewer pollen grains and deposited 2.1x fewer grains on stigmas per visit. Mobile stamens exported their pollen to significantly more neighboring flowers.

**Graphic abstract:** Behaviour and pollen transfer after flower visitors received a beating on the tongue or in the face by the forward-snapping stamens of *Berberis*. Stamens only snap forward if their filament basis is touched by an insect tongue.

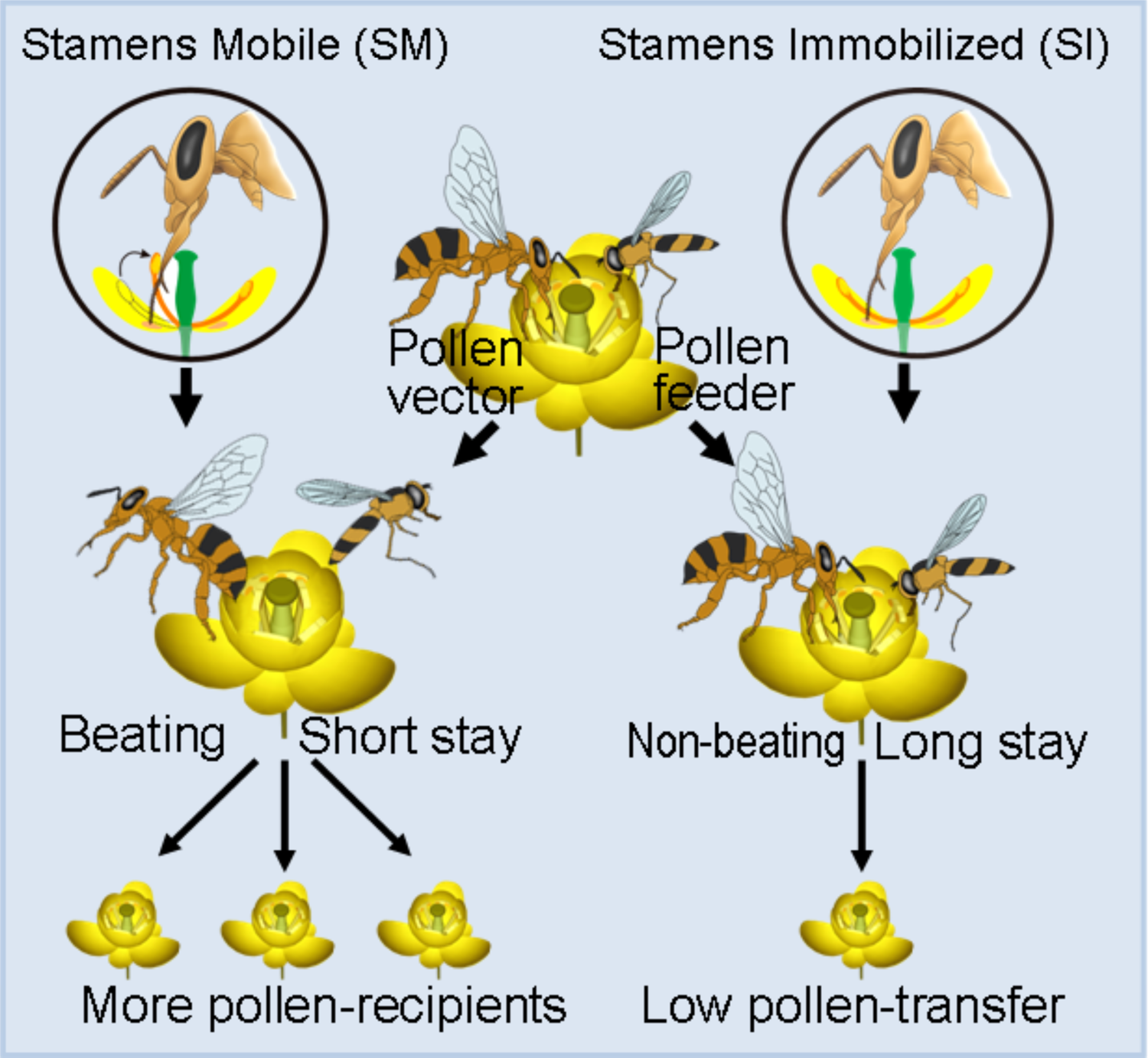

## Introduction

Animals visit flowers to forage for food or other rewards, mainly for nectar and/or pollen (Ollerton 2021). From the perspective of male reproductive success, nectar and pollen are entirely different rewards because paternity is maximized if pollen grains from one flower are deposited on multiple conspecific recipients, rather than ending up as food, while nectar is intended as pollinator food (Westerkamp 1996; Westerkamp and Claßen-Bockhoff 2007). A plant’s success as a father can depend on its temporal deployment of pollen and on the accuracy of pollen placement on the most effective pollen vectors (Harder and Thomson 1989; Armbruster et al. 2014, and studies cited therein). Flowers are therefore under selection to ‘pay’ visitors as much as possible by nectar, which can be replenished, and to meter out their non-replenishable pollen grains by placing them on multiple high-fidelity vectors. One way in which plants achieve this, is by filtering their visitors and by encouraging visitors to move on rather than stay a long time per flower.

Biologists from Linnaeus (1755) onwards have been aware of the forward-snapping movement of the stamens of *Berberis vulgaris* once the filament basis of an individual stamen is touched by a nectar-drinking insect (or a pointed object). *Berberis* flowers have two whorls of three petals and six stamens, each inserted in the middle of a petal between two nectaries. Nectar constitutes the main floral reward. The relatively few pollen grains produced in each anther remain hidden in the paired pollen sacs until each opens by an apical valve (Fig. 1). Different species of *Berberis* vary in the rapidity of their stamen movements and also in the extent of recovery and repeatability (Percival 1965; Lechowski and Bialczyk 1992; Lebuhn and Anderson 1994; this study).

**Figure 1.**
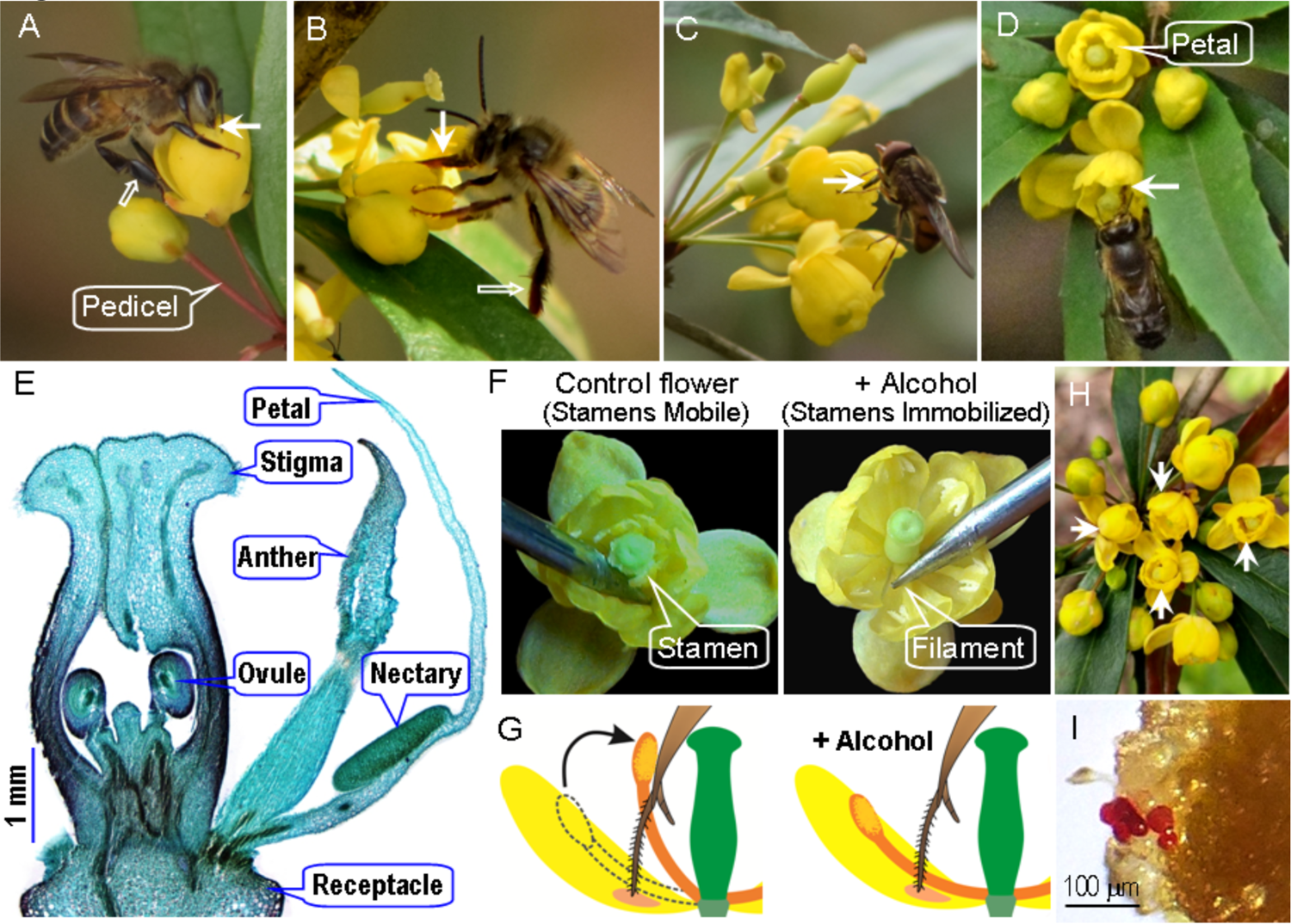
Flower traits, foraging behaviour of visitors, and manipulations of stamen movements in *Berberis julianae*, which has stamens characterized by a touch-sensitive rapid movement towards the flower centre. The major pollinators, workers of *Apis cerana* (A), and a long-tongued bee *Anthophora waltoni* (B) sucking nectar while their tongues (arrow) may contact filaments, anthers, and/or stigmas. These bees do not groom *Berberis* pollen into their corbiculae, and their legs are therefore without pollen loads (hollow arrows). Syrphid flies sucking nectar and feeding on pollen, here *Rhingia campestris* (C) functioning as a pollen thief. A bee visiting two flowers with experimentally immobilized and hence touch-insensitive stamens (D). (E) A cross section of a floral bud, showing the two anther valves and two nectaries at the base of each petal. (F) Natural flower with mobile stamens (left) snapping inward when their filament bases are touched by a needle; stamen-immobilized flower (right) whose pedicel had been immersed in 75% alcohol for over 30 min. (G) Diagram of stamen-mobile and stamen-immobilized flowers, illustrating the stamen movement when a bee’s tongue touches the filament. (H) A floral array on an inflorescence in the field with four alcohol-treated SI flowers (arrows). (I) Stained pollen grains (red) deposited on a stigma under open pollination in the field.

Early workers thought that the unidirectional stamen movements to the flower center, where the stigma is located, played a role in self-pollination (e.g., Linnaeus 1755; Sprengel 1793), but since about the 1880s, it has generally been assumed that the stamen movement helps to precisely pack pollen on the tongues or faces of flies or bees (Kirchner 1911; Knoll 1956; Percival 1965; Kugler 1970; Lebuhn and Anderson 1994). Kirchner (1911: 187-138) furthermore suggested that insects hit by a stamen would be encouraged to leave the flower, but soon would land on another flower to resume their nectaring. Rapid succession of brief visits to many flowers in Kirchner’s view should enhance cross pollination.

In this study, we experimentally test these hypotheses by immobilizing the stamens in the flowers of three species of *Berberis* and one species of *Mahonia*. The filament bending relies on rapid changes in the calcium permeability of membranes (Lechowski and Bialczyk 1992), and we therefore explored treatments with calcium inhibitors and with alcohol. We discovered that immersion of flower pedicles in 75% alcohol for 35-45 minutes was effective at blocking the stamen movement. A test for possible effects of the alcohol treatment on foraging behaviors of the major floral visitors revealed no statistical effects. We then built experimental arrays with untreated and treated flowers in enclosed conditions to quantify pollen export and import from single visits of bees and flies (set-up shown in Fig. S1). These experiments allowed us to answer the following questions (1) Are *Berberis* flowers with mobile stamens visited by the same types of insects and at the same rates as flowers with immobilized stamens? (2) Do flowers with mobile stamens have higher maternal and/or paternal success than those with immobilized stamens? (3) Do mobile stamens contribute to pollinator filtering by eliciting different behaviors in different insect taxa? And (4) is berberine, an alkaloid with antifeedant activity against herbivores that is found in *Berberis* leaves (Schmeller et al. 1997; Manosalva et al. 2019), also present in *Berberis* pollen or nectar?

## Materials and methods

### (a) Plant and insect study species

During each March between 2019 and 2022, we studied a natural population of *Berberis julianae* C.K.Schneider in a field located at 29°52′26″N, 105°30′32″E, 427 m above sea level, about 30 miles southeast of Anyue County, Sichuan Province, China. Experiments to evaluate the effect of stamen movements on pollen dispersal were also carried out in a natural population of *B. jamesiana* Forrest & W.W.Smith at Shangri-La Alpine Botanical Garden (N 27°54′05″, E 99°38′17″, 3300-3350 m above sea level), Yunnan Province, Southwestern China. Lastly, we used planted populations of *B. forrestii* Ahrendt at the Shangri-La field station and of *Mahonia bealei* (Fortune) Carrière in the Wuhan Botanical Garden (N 30°33′2″, E 114°25′48″, 23 m above sea level) in Hubei Province, China, to test the effects of the alcohol treatment on stamen mobility. Our *Berberis* and *Mahonia* taxonomy follows the Flora of China (Ying et al. 2011). Herbarium vouchers of each species have been deposited in the herbarium of Central China Normal University (CCNU). All species are hermaphroditic perennial shrubs with clusters of 9-25 yellow flowers begin opening in March or April. Individual flowers last for three to four days.

Individual insect visitors were captured with insect nets and were directly needled in the field for later identification in the last author’s laboratory in Wuhan.

### (b) Touch-sensitive stamen movements

In all our study species (as in most Berberidaceae), each anther dehisces upward by two valves exposing the pollen grains (Fig. 1). The bottle-shaped pistils have one ovary containing 2-4 ovules with a discoid stigma with a peripheral ring of papillae (Fig. 1E).

### (c) Alcohol as an inhibitor of the stamen movement and tests for confounding effects of the alcohol treatment on visitor behavior

When floral pedicels were immersed in a solution of 75% alcohol for 40 min, this blocked all stamen movement (Video 1 of *B. julianae* and video 2 of *Mahonia bealei*). To test whether any lingering alcohol scent affected visitor behavior in alcohol-treated flowers, we set up arrays with different types of flowers as follows. We first bagged >20 flowers on different individuals of *B. julianae* before they opened. Once open, 18 flowers were gently cut off and subjected to one of the three treatments: (1) stamen-mobile (**SM**) flowers: six flowers without any treatment; (2) stamen immobilized (**SI**) flowers: six flowers whose pedicels (ca. 10 mm long) were immersed in 75% alcohol for about 40 minutes; and (3) six natural flowers in a fixed position above alcohol (**SMA** flowers). For this, the pedicels of freshly opened flowers were inserted into 30 mm-long Eppendorf microcentrifuge tubes that contained 10 µL alcohol, such that they did not touch the alcohol (Fig. S1A). All 18 flowers were then fixed in position by inserting them into small holes on the surface of a paper box covered by a clean glass cup (as shown in Fig. S1). We then placed freshly caught *Berberis* visitors inside the glass cup so they could interact with the enclosed floral arrays and recorded visitation rates (visits per flower per 10 min) and handling times.

To further examine any possible effects of the alcohol treatment versus the loss of stamen movements on the duration of insect visits, we set up additional floral arrays consisting of three types of flowers: six SM flowers, six alcohol-treated SI flowers, and six filament-damaged (**FD**) flowers, the latter being flowers in which all six filaments were damaged with clean forceps so that the stamens could not move but petal nectaries and the anther sacs were still there (Fig. S1C). The experimental procedure for these arrays was the same as above (Fig. S1B).

### (d) The effect of stamen snapping on visitor behavior and pollen export and import

Flowers of *B. julianae* were mainly visited by two species of bees and two species of syrphid flies (*Results*). To examine effects of stamen snapping on foraging behavior and pollen transfer, floral visitors were allowed to interact with the above-described flower arrays both in the field (Fig. 1) and under enclosed conditions (Fig. S1D). Under enclosed conditions, we compared visitation rates, handling times, numbers of stamens touched, nectar volume remaining, pollen removal, and pollen deposition on the stigma after single visits of the four species of visitors. Sample sizes, observed trials, and measured parameters are presented in Supplementary Method S(d).

### (e) Fruit and seed set after self-pollination vs. cross-pollination, and pollen export/import after single visits

To test whether *B. julianae* is self-compatible and fruit/seed production is pollinator-limited, in 2020 and 2021, 80 flowers from 12 individuals were subjected to the following four pollination treatments. (1) Control: 20 randomly chosen flowers from 12 individuals were open pollinated, while the remaining 60 flowers were bagged throughout and subject to one of three treatments: (a) automatic self-pollination: flowers not manipulated, (b) self-pollination: flowers hand-pollinated with pollen mixtures collected from flowers of the same individual, (c) cross-pollination: flowers hand-pollinated with pollen mixtures from multiple flowers of individuals at least 20 m away. Flowers were then bagged with mesh until the petals dropped one week later. Fruits were harvested three months later and seeds and undeveloped ovules per fruit counted. Fruit set was calculated as fruit number divided by the total flower number in each treatment. Seed set was calculated as seed number per fruit divided by total number of seeds and undeveloped ovules. Aborted fruits were not included.

To examine the capability of intra-flower self-pollination induced by the stamen movement, we counted pollen grains deposited per stigma under a light microscope in 16 bagged flowers in which we had triggered one stamen by a needle. To quantify the pollen loads placed on insect bodies and the stigmas of next-visited conspecific flowers, we made an individual bee’s tongue contact all six filaments of each flower to simulate bee foraging behavior on an SI or an SM flower. To estimate stigmatic pollen loads, we allowed an individual bee to visit three trials with SM and SI flowers: Trial (1) SM + SM, Trial (2) SM + SI, and Trial (3) SI + SI flowers, see Supplementary Method S(e).

### (f) Effects of stamen snapping on pollen flow

To compare paternal success between SM and SI flowers, we conducted four pollen tracking experiments each in *B. julianae* in 2022 and *B. jamesiana* in 2021. Each trial was conducted on a fine day and involved 60 flowers from 3-5 individuals whose pollen was stained as the pollen donor: 30 flowers were alcohol-treated (SI flowers) and the remaining 30 flowers were SM flowers. Dyed pollen grains deposited on each stigma of >100 flowers (126-260) from nearby racemes of the same or different individuals were counted and the straight-line distance (within 50 cm and over 50 cm) from the racemes with the receiving flowers to the racemes with donor SM or SI flowers was recorded with a meter rule, see Supplementary Method S(f).

### (g) Statistical analyses

Visitation rates of visitor species in the field (not normally distributed) were examined with nonparametric Kruskal-Wallis tests. To compare visitor behavior on SM, SMA or FD, and SI flowers, we performed a GLM analysis with normal distributions and identity-link function to test for differences in visitation rates, handling time, and pollen transfer efficiency (log10-transformed) of the four main visitor species. Nectar volumes remaining after different visitors had visited were examined with a nonparametric Kruskal-Wallis test. To compare the number of stamens touched in different flowers, pollen removal, and pollen deposition by each visitor species, as well as pollen export and import, we performed GLM analyses with Poisson distributions and loglinear-link function. A goodness of fit-test (McDonald 2009) was used to examine whether pollen dispersal from SM and SI donors differs from a 50:50 null model at short and long distances. A GLM with binomial distribution and logistic-link function was used to detect the effects of the selfing and outcrossing on fruit set and seed set (with fruit/seed number as event variable, total treated flower/ovule number as trial variable, and different treatments as factors). The GLM analyses were performed in SPSS 19.0 (IBM, Armonk, NY, USA), and the G-test was performed in the package DescTools in R V.3.5.0 (R Core Team 2018).

## Results

### (a) Touch-sensitive stamen movements and experimental selfing and outcrossing

Flowers of *Berberis* and *Mahonia* have six petals, each with two basal nectar glands, and six stamens inserted between these glands (organ sizes are given in Table S1; videos 1 and 2). Each pollen sac opens by a separate valve and contains about 610 ± 6 grains sticky yellow pollen grains that remain attached to the pollen sac (Fig. S2G). When a flower visitor (or a pointed object) contacts the basis of a filament, the respective stamen snaps forward (towards the flower center), placing pollen grains on the visitors’ tongues or faces (Fig. 1F; Fig. S2E and F), usually on the tongue (proboscis) if the insect has been drinking nectar (Fig. 1A-C). The stamen movement takes 0.44 ± 0.02 s in *B. julianae*, 0.17 ± 0.02 s in *B. jamesiana*, and 0.23 ± 0.04 s in *B. forrestii* (Table S1). Within one minute, the stamen moves back from the flower centre, where the single style with its large stigma is located, to the petal, taking 227.70 ± 10.06 s, 110.37 ± 6.64 s, and 155.31 ± 14.07 s, respectively, to return its original position (Table S1). In *Mahonia bealei*, the stamen movement takes 0.10 ± 0.01 s (N = 10 flowers), and 3.46 ± 0.71 s later, the stamen starts moving back, taking 7.74 ± 1.96 s to return (video 2).

In *Berberis julianae*, fruit set in open-pollinated flowers was significantly higher (80.0% ± 6.4%) than that in flowers bagged to exclude pollinators (27.5% ± 7.1%), which were self-fertilized (Fig. S4). However, seed set per fruit did not differ between selfed and cross-pollinated flowers.

### (b) Flower visitors and pollinators

At our study site, B*. julianae* was visited mainly by four insect species (Table S2), the bees *Apis cerana* Fab., 1793 (Fig. 1A) and *Anthophora waltoni* Cockerell, 1910 (Fig. 1B) and the flies *Rhingia campestris* Meigen, 1822 and *Sphaerophoria menthastri* (Linnaeus, 1858). The bees foraged for nectar, but not pollen, while the flies fed on both nectar and pollen (Fig. 1C). Consistent with these feeding habits, pollen transfer efficiency of the bees was significantly higher than that of the flies (Table S2).

### (c) Tests for a possible confounding effect of the alcohol treatment on visitor behavior

We found no effect of any lingering alcohol scents in stamen-immobilized flowers on visitor behavior. Visitation rates of *Apis cerana* to *B. julianae* under enclosed conditions did not differ between untreated flowers with mobile stamens (SM), untreated flowers in a fixed position above alcohol (SMA), and SI flowers with immobilized stamens via alcohol immersion (Wald χ^2^ = 0.194, df = 2, P = 0.908; Fig. 2A). However, *A. cerana* stayed longer in in SI compared to SM and SMA flowers (Wald χ^2^ = 64.599, df = 2, P < 0.001; Fig. 2B), showing that it was the stamen snapping that caused these bees to leave flowers.

**Figure 2.**
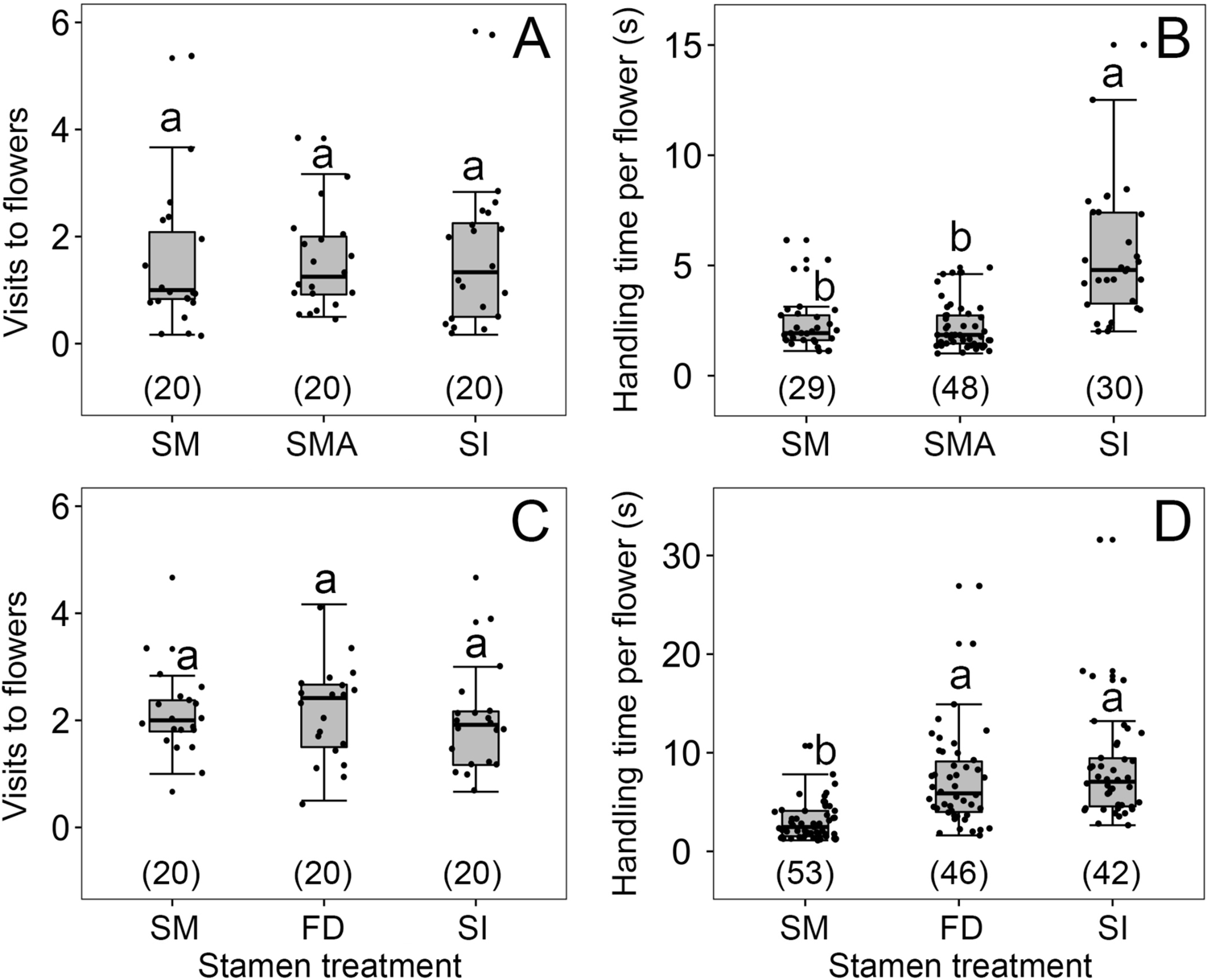
Visitation rates (A, C) and handling times (B, D) of *Apis cerana* in four types of flowers of *Berberis julianae*: Stamens mobile (SM, controls), stamens immobilized (SI), natural flowers in a fixed position above alcohol (SMA), and flowers with their filaments damaged (FD) so that the stamens became immobile but retained their pollen sacs and the nectar glands to the right and left of each filament. The box plots indicate median (mid lines), inter quartile range (boxes) and 1.5 times the inter quartile range (whiskers) as a well as outliers (points). Different lowercase letters indicate significant differences, and numbers in brackets indicate sample sizes.

Visitation rates of *A. cerana* to *B. julianae* under enclosed conditions also did not differ among SM, SI, and filament-damaged (FD) flowers (Wald χ^2^ = 0.44, df = 2, P = 0.802; Fig. 2C), and bees stayed less time in SM flowers than in flowers without stamen snapping (SI and FD flowers), but equally long in SI and FD flowers, showing that it was the stamen snapping rather than any lingering scent from the alcohol treatment that caused visitors to move on.

### (d) Effect of stamen snapping on visitor behavior and pollination in the field and under enclosed conditions

In the field, when three or four SI flowers of *B. julianae* were attached to natural inflorescences with the same number of SM flowers (Fig. 1H), *Apis cerana* stayed much longer on SI flowers than on SM flowers (15.46 ± 1.54 s vs. 3.63 ± 0.33 s, Wald χ^2^ = 56.055, P < 0.001 in 2020; 16.076 ± 1.515 s vs. 5.675 ± 0.382 s, Wald χ^2^ = 68.421, P < 0.001 in 2021). Despite the longer visits, many fewer pollen grains were loaded onto *A. cerana* after a single visit to SI flowers than to SM flowers (716 ± 85 vs. 1223 ± 100 grains in 2020; 890 ± 73 vs. 1401 ± 134 grains in 2021; Wald χ^2^ = 12.873, P < 0.001 in 2020; Wald χ^2^ = 10.511, P = 0.001 in 2021), and the numbers of pollen grains deposited on stigmas by a single *A. cerana* visit also were much lower in SI flowers than in SM flowers (87 ± 10 vs. 230 ± 25 grains in 2020; 63 ± 8 vs. 260 ± 35 grains in 2021; Wald χ^2^ = 26.847, P < 0.001 in 2020; Wald χ^2^ = 40.042, P < 0.001 in 2021; Fig. S5). Pollen transfer efficiency by *A. cerana* was therefore reduced in SI compared to SM flowers, and this was significant in 2021 but not 2020 (0.084 ± 0.016 vs. 0.303 ± 0.102; Wald χ^2^ = 4.505, P = 0.034 in 2021; 0.205 ± 0.024 vs. 0.246 ± 0.091; Waldχ^2^ = 0.188, P = 0.665 in 2020; Fig. S5 I-J).

When the experiment was repeated in 2021 with five SM and five SI flowers under enclosed condition (Fig. S1), both bee species and both syrphid fly species stayed longer on SI than on SM flowers (33.7 ± 4.2 s vs. 16.0 ± 1.7 s; Wald χ^2^ = 30.106, P < 0.001; Fig. 3E′-T; Table S3). All four visitor types exploited more nectar (Fig. 3M′-P) and touched more stamens in SI than in SM flowers (Fig. 3I’-L). Pollen transfer efficiency did not differ between the two bees, but was higher than that of the two flies (Wald χ^2^ = 13.319, df = 3, P = 0.004, N = 80). In SI flowers, *A. cerana* was loaded with fewer grains than in SM flowers, while both *Anthophora waltoni* and the flies removed more grains from SI flowers than from SM flowers (Fig. 3Q’-T).

**Figure 3.**
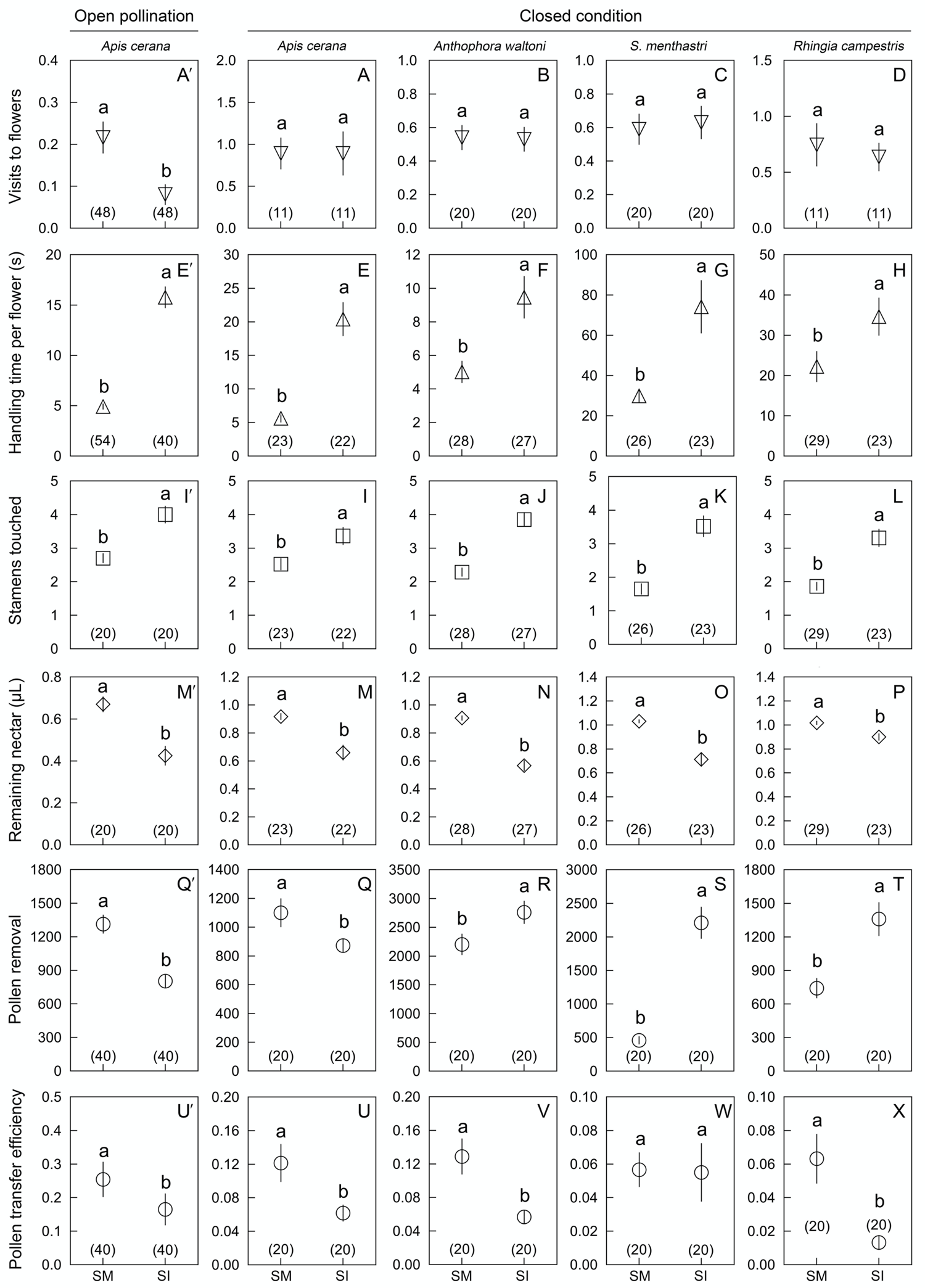
Comparisons of six parameters (mean ± standard error) in *Berberis julianae* used to examine the effects of stamen movements on insect visitor foraging behavior and their roles in pollination. The major pollinator, *Apis cerana*, was studied under both open pollination (far left) and enclosed conditions (set-up shown in Fig. S1 and video 3), whereas the infrequent bee pollinator, *Anthophora waltoni*, and the two fly pollen thieves (*Sphaerophoria menthastri* and *Rhingia campestris*) were compared under enclosed conditions. Different lowercase letters above error bars indicate significant differences between control (stamen mobile, SM) and alcohol-treated (stamen immobilized, SI) flowers. (A′-D) Visitation rates of four visitor species, showing that *A. cerana* visited control flowers more frequently than SI flowers under open pollination (A′), but no visitor species discriminated between SI and SM flowers under enclosed conditions (A-D). All visitor species spent more time (E′-H) and touched more stamens (I′-L) in SI flowers than in control SM flowers. Visitors removed more nectar from SI flowers, resulting in less nectar remaining per flower (M′-P). Pollen removal by *A. cerana* was lower from SI than from SM flowers (Q′ and Q), but higher in the other three visitor species (R-T). Compared to SM flowers, pollen transfer efficiency was significantly decreased in SI flowers (U′, U, V and X) although it did not differ in *Sphaerophoria menthastri* (W).

### (e) Effects of stamen snapping on self pollination and pollen transfer in single visits

Self-pollen receipt by stigmas after a single stamen movement (14 ± 3 grains, N = 16) was only 6% of the pollen receipt resulting from a single *A. cerana* visit (245 ± 22) or 1% of total pollen of a single anther, indicating that intra-flower self-pollination mediated by the stamen movements plays a minor role in total pollen receipt. Both bee species *A. cerana* (Fig. 4A) and *A. waltoni* (Fig. 4B), carried fewer pollen grains on their tongues after visiting SI flowers compared to SM flowers (Wald χ^2^ = 142.565, P < 0.001 and Wald χ^2^ = 14.236, P < 0.001, respectively). In three trials with alternative floral arrays (Materials and Methods section (e)), the number of pollen grains deposited on stigmas by *A. cerana* (mean ± SE, Fig. 4C) and *A. waltoni* (mean ± SE, Fig. 4D) in Trial 1 (SM + SM) was significantly higher than in Trials 2 (SM + SI) and 3 (SI + SI; Wald χ^2^ = 118.887, P < 0.001 and Wald χ^2^ = 69.274, P < 0.001, respectively). Moreover, pollen deposition by *A. cerana* or *A. waltoni* was significantly higher (P < 0.001) in Trial 2 (SM + SI) than in Trial 3 (SI + SI), indicating that stamen snapping facilitates pollen export.

**Figure 4.**
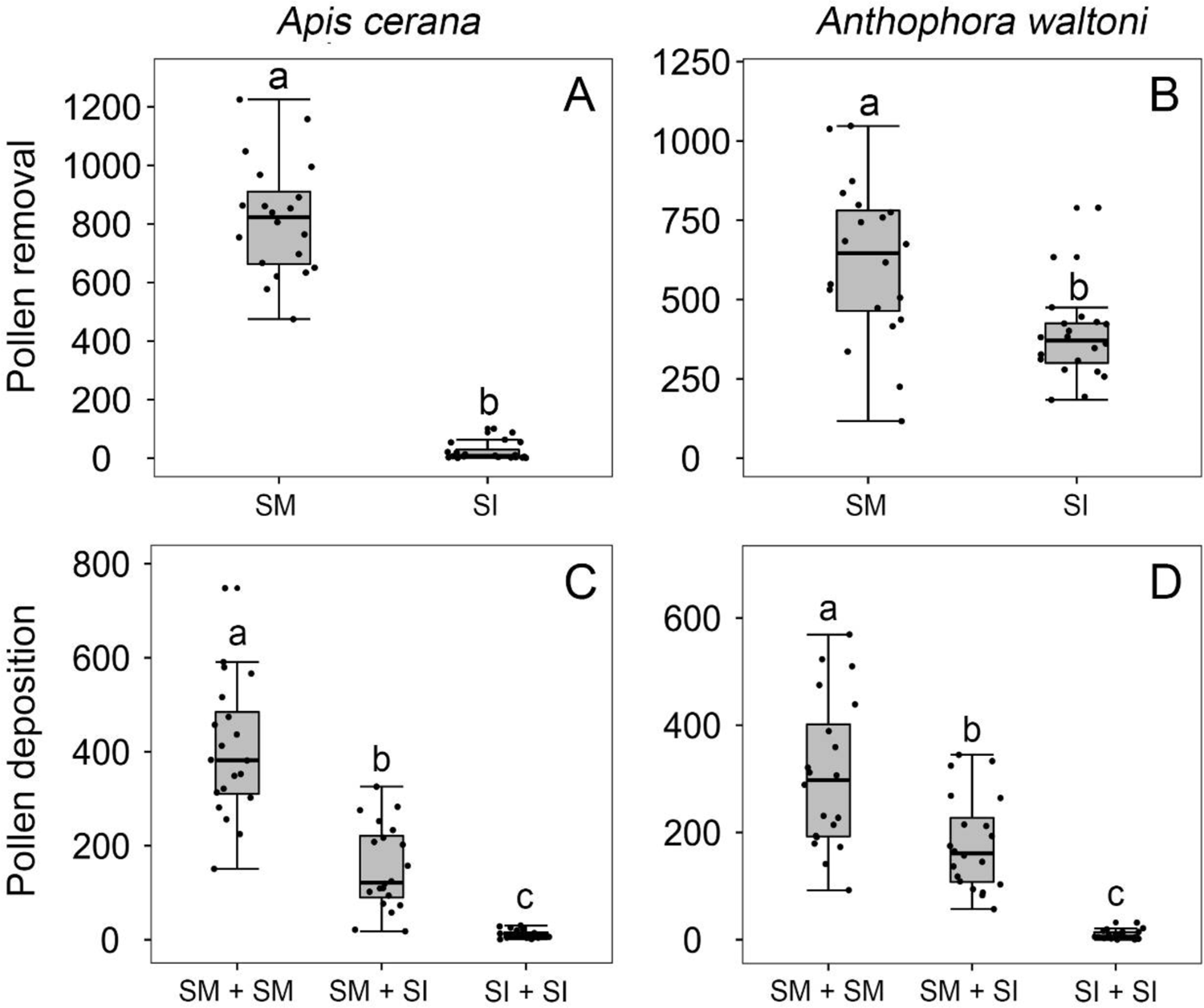
Effects of stamen snapping on pollen placement on the pollinator body and pollen deposition on stigmas after single visits by *Apis cerana* (A, C) and *Anthophora waltoni* (B, D). Numbers of pollen grains placed on bees’ tongues during a single visit were significantly higher when stamens were mobile (SM) than when stamens were experimentally immobilized (SI) (A, B). Numbers of pollen grains deposited on the stigma of the second-visited flower (pollen recipient) in a single visit by *Apis cerana* (C) and *Anthophora waltoni* (D) in three trials with experimentally treated and control flowers (SM + SM: the bee visiting two SM flowers; SM + SI: first visiting one SM and then one SI flower; and SI + SI: visiting two SI flowers. The box plots indicate median (mid lines), inter quartile range (boxes) and 1.5 times the inter quartile range (whiskers) as a well as outliers (points). Different lowercase letters indicate significant differences among three trials.

### (f) Effects of stamen snapping on paternal success

Paternal success of SM and SI flowers was quantified in our pollen tracking experiments in which pollen of flowers of *B. julianae* and *B. jamesiana* was stained *in situ* with either eosin or aniline blue and stigmas of all flowers in the vicinity were then checked for the stained grains.

Of over 700 flowers of *B. julianae* whose stigmas we checked, 44 received pollen from SM flowers and 14 from SI flowers, indicating that flowers with mobile stamens donated pollen to about 3.1x more flowers (44/772 vs 14/733, G = 38.845, p < 0.001, Table S4). The mean number of pollen grains deposited per stigma was also higher in SM than SI flowers (0.16 ± 0.03 vs. 0.03 ± 0.01; Wald χ^2^ = 76.536, *P* < 0.001). All four runs of this experiment showed a consistent pattern: more pollen grains from SM flowers were delivered to more flowers (Fig. 5), while SI flowers donated fewer grains to fewer recipients. Within <50 cm from the dyed pollen source, 25 of 338 flowers received pollen from SM flowers, while only 10 of 395 flowers received pollen from SI flowers (G = 21.555, p < 0.001, Table S4). At distances >50 cm, 19 of 434 flowers received pollen from SM flowers, while only 4 of 338 flowers received pollen from SI flowers (G = 22.435, p < 0.001, Table S4).

**Figure 5.**
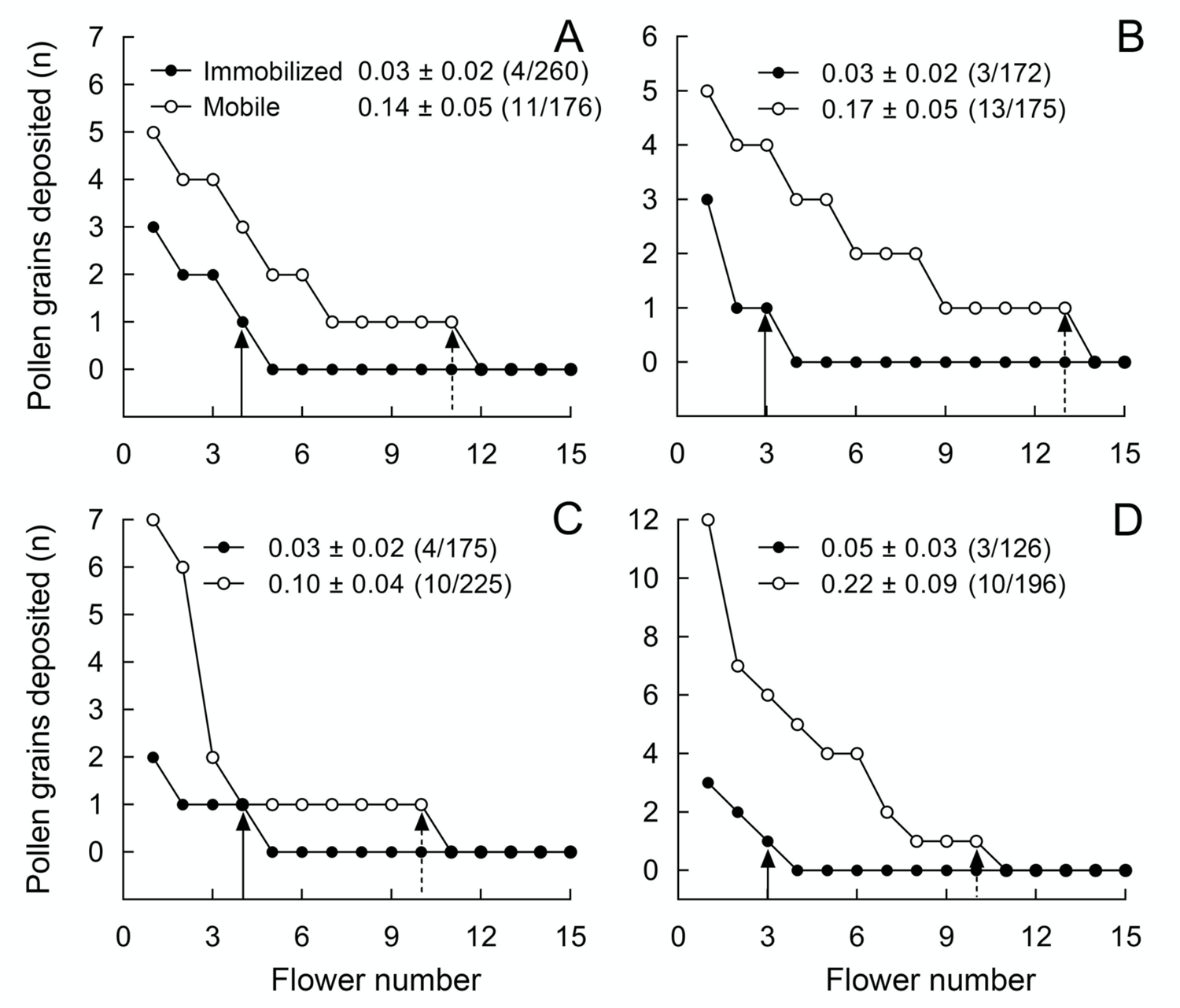
Number of stained pollen grains deposited on the stigmas of control flowers with mobile stamens (open circles) and on stigmas of flowers with experimentally immobilized stamens (closed circles) in *Berberis julianae*. Mean pollen number and standard errors (Numbers of flowers with stained pollen deposition/total number of sampled flowers of pollen recipients) are given for treated and control flowers in each of four trials (A, B, C and D). Note that only 15 flowers are shown, although each pollen-tracking test sampled over 100 flowers to examine the effect of stamen movements on pollen dispersal; for example in 260 pollen-recipient flowers, only four flowers had received pollen grains from stained SI flowers, i.e., (4/260) in (A).

Of 400 nearby flowers in *B. jamesiana,* 75 received pollen from SM flowers and 40 from SI flowers, again indicating that mobile stamens donated pollen to more flowers (75/400 vs 40/400, G = 27.81, p < 0.001). The mean number of pollen grains deposited per stigma was also higher in SM than SI flowers (0.4 ± 0.05 vs. 0.2 ± 0.04; Wald χ^2^ = 22.95, P < 0.001). All four runs of this experiment showed the same pattern as found in *B. julianae*. Within <25 cm from the dyed pollen source, 13 of 60 flowers received pollen from SM flowers, while 9 of 46 flowers received pollen from SI flowers (G = 0.164, p = 0.685). At distances >25 cm, 7 of 60 flowers received pollen from SM donors, while only 1 of 56 flowers received pollen from SI flowers G = 15.037, p = 0.0001, Table S5).

### (g) Berberine content in *B*. *julianae* leaves, petals, pollen, and nectar

High-performance liquid chromatography of the berberine concentrations in *B. julianae* leaves, petals, pollen, and nectar indicated high berberine concentrations in leaves, petals, and pollen, while no berberine was detectable in the nectar (Table S6). That bees can taste the berberine in the pollen is suggested from the observation that individuals of both bee species used their front legs to remove pollen grains that stuck to their tongues.

## Discussion

To maximize pollen dispersal, the number of grains removed by each visitor should be limited so as to heighten the probability of pollen reaching different conspecific stigmas (Harder and Thomson, 1989), and where possible, visitors should be ‘paid’ by replenishable nectar rather than pollen (Westerkamp 1996). Especially in flowers with scarce pollen grains, such as *Berberis* and *Mahonia*, it should also be advantageous to protect pollen grains from exploitation by pollen thieves and inefficient vectors by physical or chemical mechanisms of defense (Palmer-Young et al. 2019; Xiong et al. 2019; Wang et al. 2019). The yellow pollen grains of *Berberis* and *Mahonia*, presented on the exposed yellow anthers without physical protection, indeed may have some chemical defense via their berberine content (Table S6), an alkaloid with antifeedant activity against herbivores and pests (Schmeller et al. 1997; Manosalva et al. 2019). In *B. julianae*, we often observed pollen on bees’ tongues or faces, but it was never groomed into the corbiculae (Figs. 1 & S2). Instead, *A. cerana* and *A. waltoni* bees cleaned off and discarded pollen grains that stuck to their tongues with their front legs. The syrphid flies, by contrast, fed on the pollen grains despite their berberine content (Table S2).

In stamen-snapping flowers, both types of bees, *Apis cerana* and *Anthophora waltoni*, typically stayed for five seconds and triggered 2-3 stamens per flower per visit (Fig. 3), a similar number as in *B. thunbergii* in North America, where the main visitors were medium-size bees of *Apis mellifera* and Anthophoridae (Lebuhn & Anderson 1994). In stamen-immobilized (SI) flowers, bees stayed about 3x longer (on average 14.37 ± 1.53) and flies about 2x longer (on average 54.38 ± 7.53 seconds), and both visitor types therefore removed more nectar. The impact of longer stays on pollen removal, however, differed between the most common bee visitor, *A. cerana*, and the other three visitors: When immobilized stamens no longer smeared pollen grains on its tongue or face, *A. cerana* carried away many fewer grains, while the pollen-feeding syrphid flies and the long-tongued bee *Anthophora waltoni* removed more grains, in the case of the flies because they ate more pollen and in the case of the long-tongued bees because they passively touched more untriggered open anther valves, the longer they stayed.

Although less efficient pollen vectors, the pollen-feeding syrphid flies visiting *Berberis* still contributed to pollination (Table S1) and were only conditional pollen thieves (Hargreaves et al., 2009). The main reason for this is that *B. julianae* is self-compatible, with seed set per fruit not differing between selfed and cross-pollinated flowers so that even the 6% self-pollen deposited on stigmas by stamens hitting the flower’s own stigma contribute to reproductive insurance. We also did not find a difference between flies and bees in their reaction to the snapping of stamens: Both visitor types left flowers more quickly after being hit by stamens, and stamen snapping therefore did not result in visitor filtering.

The quicker leaving of flowers greatly increased male reproductive success by dispensing pollen to multiple recipients. This is evident from pollen dispersal distances in SM and SI flowers of *B. julianae* (Fig. 5) and *B. jamesiana* (Fig. S6 and Table S5), and from the proportions of pollen recipients that were reduced by 60.0% and 36.4% in the two species. Studies on other species with stamens triggered by flower visitors have proposed that touch-sensitive stamens may discourage pollen feeders (Schlindwein & Wittman 1997: generalist bees in *Opuntia brunneogemmia*; Wong Sato & Kato 2018: pollen-harvesting bumblebees in *Meliosma tenuis*). Our experiments in *Berberis* support this hypothesis although the situation is different because *Berberis* pollen appears to be unattractive to bees due to its chemical composition, while in both these studies bees actively collected the pollen.

A final function of the mobile stamens in *Berberis* is that they enhance pollination accuracy in the bee visitors because of the accurate positioning of pollen on a place of the bees’ bodies that would contact the stigma of the next-visited conspecific flower.

## Conclusion

Even though botanists have speculated about the adaptive value of the visitor-triggered forcefully snapping stamens of *Berberis* since 1755 (Linnaeus 1755), this is the first experimental investigation of how this trait impacts the flowers’ paternal and maternal fitness. Our results demonstrate surprisingly large effects and reveal another mechanism by which plants simultaneously meter out their pollen and reduce pollen theft.

## Authors’ Contributions

S.-Q.H. and D.-F.L. designed the research; D.-F.L., and W.-L.H. performed the research; and S.-Q.H. assisted the field research. S.-Q.H., D.-F.L. and S.S.R. analyzed the data and wrote the paper. All authors commented on the manuscript.

## Acknowledgments

We thank lab members Q.-M. Quan, X.-W. Lv and Z.-X. Tian for field assistance and Z.-Y. Tong, and Y.-Z. Xiong for methodological and statistical advice; Professor H.-L. Xu for identifying insects; Dr. Chih-Chieh Yu for confirming our plant identifications; Z.-D. Fang and staff of Shangri-La Botanical Garden for logistical support; and Sarah Corbet, Steven Johnson, and Nathan Muchhala for sharing ideas and providing helpful suggestions on early version of the manuscript.

## Supplemental materials: Detailed descriptions of methods and 6 tables, 6 figures, and 3 videos, the latter uploaded separately

### Supplemental Methods

#### (a) Measurements of floral traits in *Berberis* and *Mahonia* species

To compare floral traits among species, flower size, length of the pedicel and other related traits were also measured with a digital caliper to 0.01 mm in 20 flowers per species (Table S1). To estimate nectar volume per flower of *B. julianae*, clean 10 µL microliter syringes (Agilent Technologies Inc., USA) were used to extract nectar drops from bagged flowers (Wang et al. 2019). To estimate the production of pollen grains and ovules per flower, we randomly collected one of six anthers from virgin flowers that had been bagged as buds with fine-mesh cotton bags to exclude visitors. Each selected anther was placed on a microscope slide and then squashed under a coverslip (Wang et al. 2019). All pollen grains were counted under a microscope and the number in one anther was multiplied by six to obtain the total pollen grain number per flower. Meanwhile, ovules of sampled flowers were also counted. Numbers of sampled flowers are presented in Table S1.

Floral traits among species were compared under a generalized linear model (GLM) with normal distribution and identity-link function except pollen number and ovule number per flower with Poisson distribution and loglinear-link function (Table S1). Data of pistil height and stamen length as well as handling time and pollen transfer efficiency were log10-transformed to achieve normal distribution.

#### (b) Alcohol-treated time to block the stamen movement

To find the minimum time required to completely block stamen movements in *Berberis* species, we checked the response of stamen movements in a time-series of alcohol-treated flowers. In *B. julianae*, the pedicels of 64 flowers were gently cut off and the bases (about 10 mm long) were immersed in 75% alcohol. Every five minutes, 8 flowers were checked by touching the filaments of each flower with a dissecting needle to identify whether the stamens remain mobile. As above, 100 flowers of *B. jamesiana* and 80 flowers of *B. forrestii* experienced alcohol treatment, which two species were sampled in May and June 2021 at a field station, Shangri-La Alpine Botanical Garden (N 27°54′05″, E 99°38′17″, 3300-3350 m above sea level), Yunnan Province (YN), Southwestern China. Every five minutes, 10 flowers of each species were tested to see whether the stamens remained touch-sensitive (Fig. S3).

#### (c) Visitation of floral visitors

To quantify visitation rates (visits/flower/hour) of different visitors to *B. julianae*, we observed visitors in the field population of SC in March 2019, 2020 and 2021. Sessions for visitor observation in a patch containing one or two individuals with over 100 open flowers each lasted 1 hour, during 9:00-12:00 am and 12:30-16:30 pm on sunny days. The foraging behaviour of each floral visitor was recorded, particularly whether they were female and fed or groomed pollen grains into pollen loads (Wang et al. 2019). The numbers of flowers visited by any visitor in one session were recorded. At the end of an observation period, all open flowers in the patch were counted. A total of 20h, 20h and 35h of observations were obtained in the three consecutive years (Table S2).

#### (d) Effects of stamen snapping on visitor behaviour and pollen export and import

To see whether floral visitors discriminate between SI and SM flowers, we compared visitation rates of the honeybee *A. cerana* under open pollination and of the four insect species under enclosed conditions (illustrated in Fig. S1).

In the SC field population under open pollination, 72 flowers from different individuals were randomly bagged with mesh nets (Wang et al. 2019) before they opened in March 2020. Twelve SI and 12 SM flowers were observed each day. When flowers were beginning to open, we gently cut off 36 flowers at the base of the pedicels and immersed the pedicels in 75% alcohol. All stamens in each alcohol-treated flower became touch-insensitive in 40 minutes. These SI flowers were set on flowering individuals with the other 36 SM flowers as controls, allowing the honeybee to visit in open pollination conditions (Fig. 1D, H). The number of flowers visited per hour was recorded during 09:00-12:00 am and 13:00-17:00 pm for three days. Visitation rates were calculated as the number of flowers visited per hour divided by the total number of observed flowers, i.e., visits per flower per hour. In March 2021, total of 160 floral buds from five individuals were randomly bagged. As above, 80 newly opened flowers were alcohol-treated and the remaining 80 flowers were free of alcohol as SM flowers. On each observation day, we set four arrays each having four SM and four SI flowers. The visitation rates were recorded during 09:30-12:30 am and 13:00-17:00 pm for five days each with 16 SI and 16 SM flowers.

To compare the handling time, pollen removal and pollen transfer efficiency of the pollinator *A. cerana* in SM with SI flowers **under open conditions**, a total of 100 flowers from different individuals of *B. julianae* were randomly bagged with mesh before the flowers opened in March 2020 and 2021. When the flowers were newly opened, 12 flowers per day were alcohol-treated on five days per year with fine weather. Four of these SI flowers were put into each of three racemes with no open flowers (4 flowers × 3 racemes × 5 days = 60), and four bagged SM flowers from two racemes were uncovered per day (4 flowers × 2 racemes × 5 days = 40) for the honeybee visits. In the five arrays each of four flowers per day, we recorded handling time of the honeybee during each visit to one flower. To estimate pollen transfer efficiency, each year total of 20 SI and 20 SM flowers visited once by the honeybee were harvested immediately in the field. Six anthers and the stigma of each flower were collected. We counted pollen grains remaining within anthers per flower and deposited per stigma respectively under a light microscope. Pollen removal per flower was calculated as the mean total number of pollen grains (see above) minus the remaining grains. Pollen transfer efficiency was calculated as pollen deposition divided by pollen removal. In both years we obtained 20 pairs of pollen removal and deposition data for SM flowers, and 20 pairs for SI flowers. To compare the number of stamens touched and the remaining nectar volume left by the pollinator *A. cerana* in SM and SI flowers under open conditions, we recorded the number of stamens touched by the pollinator when it visited each SM or SI flower once in March 2021. Nectaries are located on both sides of each of the six stamens. In SM flowers, the filaments rapidly propel the anthers toward the pistil when touched by visitors probing for nectar. In SI flowers, when a visitor is collecting nectar from drops at the petal base, its tongue touches the filament nearby but does not trigger stamen movement. As the visitor switches to collect nectar from another drop, its body turns. Therefore, counting the visitor’s body turns allowed us to record the number of stamens touched by a visitor in SI flowers (Video 3). To estimate nectar collection by different visitors, we measured the remaining nectar volume within each SM and SI flower after one visit using a clean 10 µL microliter syringe (Agilent Technologies Inc., USA).

To compare the foraging behaviour of pollinators (*A. cerana* and *A. waltoni*) and of pollen thieves (*S. menthastri* and *R. campestris*) and their roles in pollination between SM and SI flowers under enclosed conditions (Fig. S1), we set up floral arrays each consisting of five SM and five SI flowers. The flowers were fixed by inserting the pedicels into small holes on the surface of a paper box covered by a clear glass cover. Under such enclosed conditions, we allowed an individual of each of four insect species to freely interview one floral array. We recorded the visitation rates (visits per flower per 10 min) and handling time of the visitor and the number of stamens touched by the visitor in each flower, measured the remaining nectar volume left by the visitor, and assessed pollen removal and pollen deposition by the visitor after one visit to calculate pollen transfer efficiency. It took four trials of interviews over two days to yield 20 once-visited flowers for each insect species. Four individuals of each insect species were captured in the field and released within two hours after their interviews to the two types of SM and SI flowers had taken place under the glass cover.

#### (e) Pollen export/import after single visits

To estimate the effects of the stamen movement on pollen loads placed on the pollinator body and stigmatic pollen loads, we simulated visits by each of the two bee species to SI and SM flowers. More than 320 flowers from different individuals of *B. julianae* were bagged before opening in March 2021. To estimate pollen placement on the bee *A. cerana* or *A. waltoni*, we held an individual bee with forceps which was captured in the field and anaesthetized by moderate gaseous acetic ether and made its tongue contact with all six filaments of each flower to simulate its foraging behaviour on a SI or SM flower. After probing the flower, pollen grains placed on the bee’s tongue (and occasionally on the head) were pasted onto a piece of tape. The tape was attached to a slide, then pollen grains placed on the bee’s tongue in 20 SI and 20 SM flowers were counted under a light microscope (Fig. 4A-B). To estimate stigmatic pollen loads, we allowed an individual bee to visit three trials with two of alternative SM and SI flowers: Trial (1) SM + SM, the bee visiting two SM flowers; Trial (2) SM + SI, first visiting one SM and then one SI flower; Trial (3) SI + SI, visiting two SI flowers. Each trial of *A. cerana* or *A. waltoni* was repeated 20 times. After each trial, the stigma of the latter flower (pollen recipient) was collected to count pollen grains deposited (Fig. 4C-D).

#### (f) Effects of stamen movement on pollen flow

To examine the effects of stamen movements on the fate of pollen, we conducted four trials of pollen tracking experiments to compare pollen paternal success between SM and SI flowers in a transplanted population of *Berberis jamesiana* at Shangri-La Alpine Botanical Garden, where the interference of stained pollen with the sexual reproduction of wild plants could be minimal. Each trial conducted on a fine day involved 60 flowers from 3-5 individuals that pollen was stained as pollen donor source: 30 flowers were alcohol-treated (SI flowers) and the remaining 30 flowers were used as SM flowers.

Pollen grains in SM or SI flowers were stained with eosin (stained red) or aniline blue (blue) respectively. The two dyes were alternately used between SM and SI flowers in four trials. The dyes dried within 5-10 minutes, and all pollen-stained flowers were taken to the field and put into racemes of two flowering individuals along the roadside for pollinators to visit. To track pollen flow, the two clusters containing either 30 SM or SI flowers were arranged separately (about 100 cm apart) on the flowering branches.

Flowers of each cluster were within a 40 cm × 40 cm square on four erect racemes. Previous studies indicated that pollen flow mediated by generalist insects usually disperses within meters with a highly leptokurtic distribution. One day (24 h) later, given that honeybee visits were infrequent to *B. jamesiana* in May 2021, stigmas of 100 flowers from nearby racemes were collected. Dyed pollen grains deposited on each stigma were counted under a stereomicroscope.

To compare the distance of pollen dispersal from SM and SI flowers, we conducted two trials of pollen tracking experiments to compare distance of pollen dispersal between SM and SI flowers of *B. jamesiana*. Each trial was the same as above. At 17:30 p.m. on the day, stigmas of all flowers from nearby racemes were collected and we noted the straight-line distance within 25 cm and over 25 cm from the racemes with the sampled flowers to the racemes with pollen-stained SM or SI flowers. Dyed pollen grains deposited on each stigma were counted under a stereomicroscope.

#### (g) Measurement of chemical defence in pollen and nectar

To examine whether berberine is present in *Berberis* pollen and nectar, we collected three separate tissues as well as nectar from 10 plants of *B. julianae*. Leaves, petals, and pollen grains were kept dry using an oven at 65°C, while the nectar was stored at −20°C before chromatographic analysis. The berberine content of the different samples was analysed using high-performance liquid chromatography (HPLC). To extract berberine from leaves, a 0.1 g leaf sample was weighed by a balance (Sartorius BAS124S) and transferred to a 2-ml micro centrifuge tube. A steel bead was added to the tube. The leaves were ground using a tissue homogenizer (Tissuelyser, QIAGEN) at 30 Hz for 10 min, and then the leaf sample was transferred to a vial. Enzymolysis of the leaf sample was conducted by adding 2 ml pure water, 1 μl dilute sulphuric acid and 6 mg cellulose at 50°C in a magnetic stirrer for 3 h. The leaf sample was extracted for 2.5 h in 5ml pure water, and then transferred to a 5-mL micro centrifuge tube. Following centrifugation at 8 000 rpm for 10 min (Centrifuge 5430R, Eppendorf), the supernatant was transferred to a vial. The volume of the sample was recorded. Berberine in petals, nectar, pollen grains was measured similarly.

Components were separated using an Acquity HPLC BEH C18 column (50 x 2.1 mm, 1.7 μm) (Waters, Milford, MA, US) set at 30°C, and the injection volume was 20 µL. All aspects of system operation and data acquisition were controlled by software (Agligent1100 series) in the Center of Analysis and Test of Wuhan University, Wuhan, China. The mobile phase was acetonitrile-0.3% phosphoric acid-pure water (35:5:60); flow rate: 1.0mL/min; determination wavelength: 346nm; the detection signal was DAD (Diode Array Detector). The berberine concentrations in the different samples was compared using a GLM with normal distribution and identity-link function.

### Supplementary figures S1-S6

**Figure S1.**
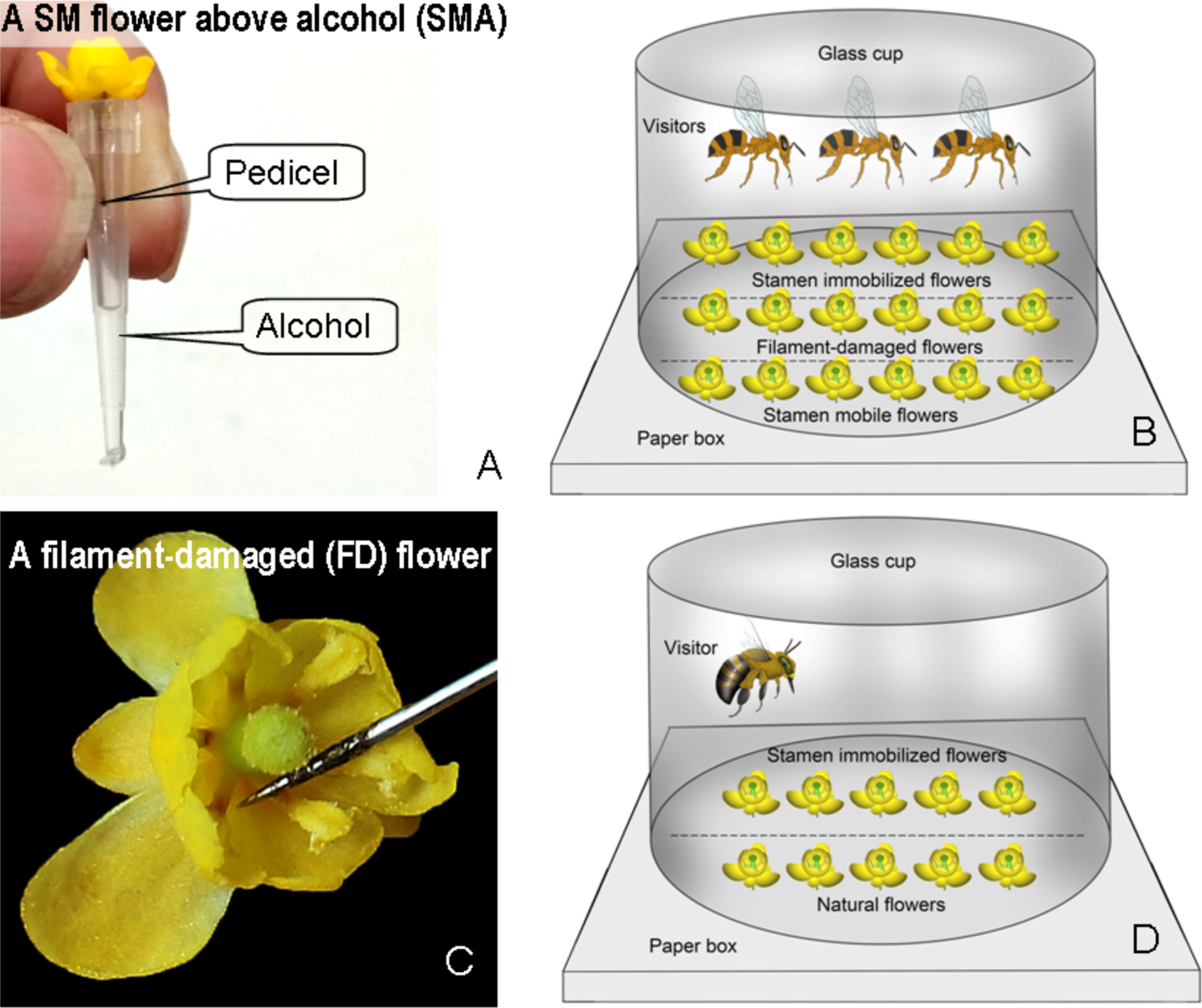
Flower manipulations and experimental floral arrays in *Berberis julianae*. (A) An SMA flower with its pedicel fixed above 10 µL of alcohol. (B) An experimental array with SM, SI, and FD flowers under enclosed conditions with three individuals of *Apis cerana*. (C) An FD flower in which all filaments have been damaged, but anthers and nectaries were intact. (D) An experimental array with SM and SI flowers under enclosed conditions with one of the four visitor species caught while foraging on flowers nearby.

**Figure S2.**
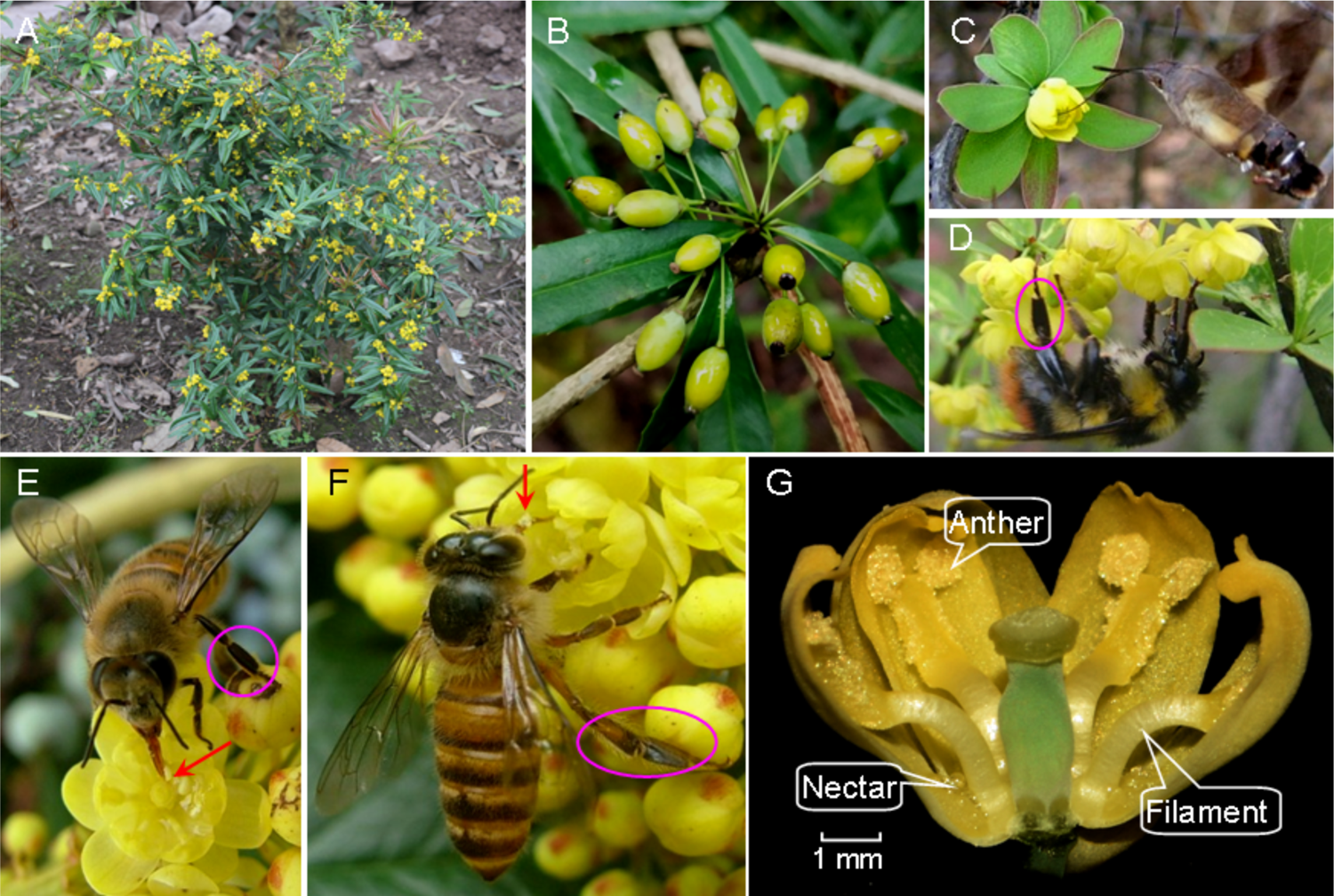
Habit, floral traits, developing berries, and feeding behaviour of various insects in three Berberidaceae taxa whose stamens are characterized by touch-sensitive rapid movements. Bird’s eye view of a flowering individual in an open habitat (A) and an infructescence (B) of *Berberis julianae* in the Sichuan population. A hawkmoth (C) and a bumblebee worker (*Bombus friseanus* Skorikov, 1933) (D) drinking nectar from *Berberis julianae*. (E) a worker of *Apis cerana* drinking nectar from *Mahonia bealei;* its tongue has triggered the anthers moving inward (red arrow). (F) Pollen grains on the tongue of *A. cerana* visiting *M. bealei*. The bees’ legs are without visible pollen loads (pink circles), showing that these bees do not actively collect *M. bealei* pollen. (G) Cross section of a *M. bealei* flower showing the two anther valves with pollen grains attached.

**Figure S3.**
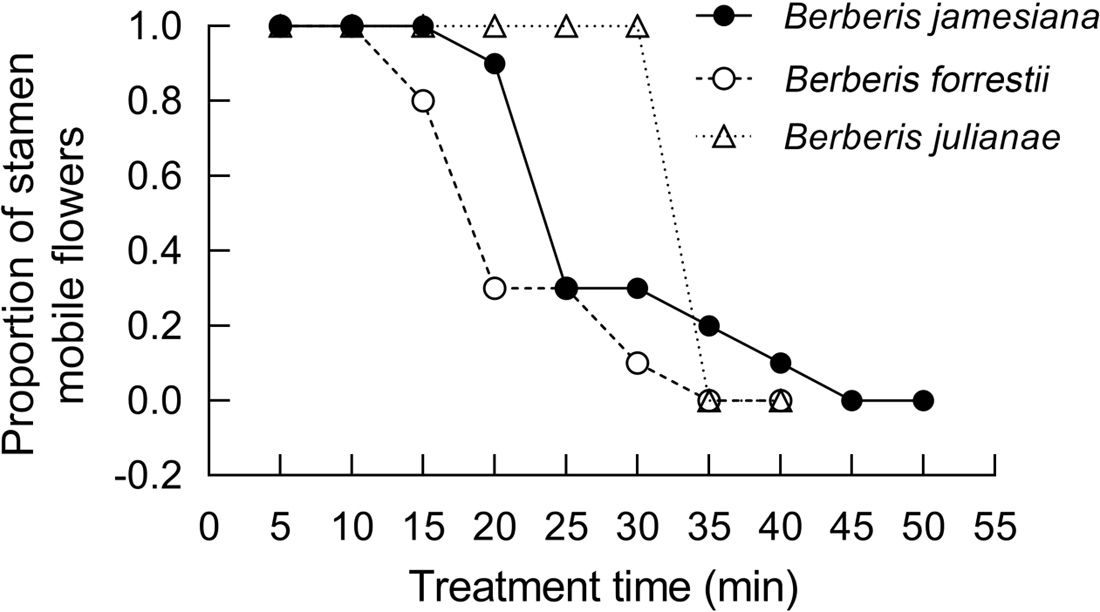
Time required for alcohol uptake to block stamen movements in three species of *Berberis*. The stamens in *B. julianae*, *B. forrestii* and *B. jamesiana* became touch-insensitive when the floral pedicels had been immersed in alcohol for 35 min, 35 min, and 45 min, respectively.

**Figure S4.**
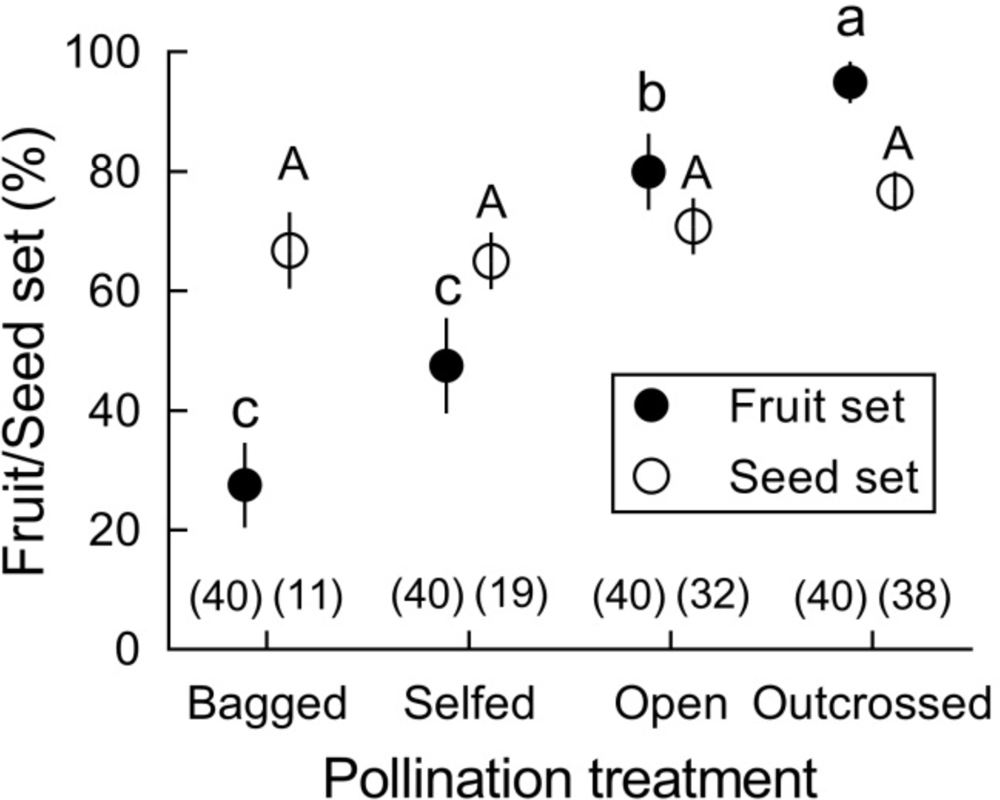
Fruit/seed set (mean ± SE) after four pollination treatments in *Berberis julianae*. Different letters beside mean values indicate significant differences among the four treatments under a generalized linear model (GLM). Fruit set differed significantly (Wald χ2 = 34.598, *P* < 0.001, df = 3) but seed set per fruit did not when zero data were excluded (Wald χ2 = 1.973, *P* = 0.578, df = 3). Sample size for each treatment is given in brackets above the X-axis.

**Figure S5.**
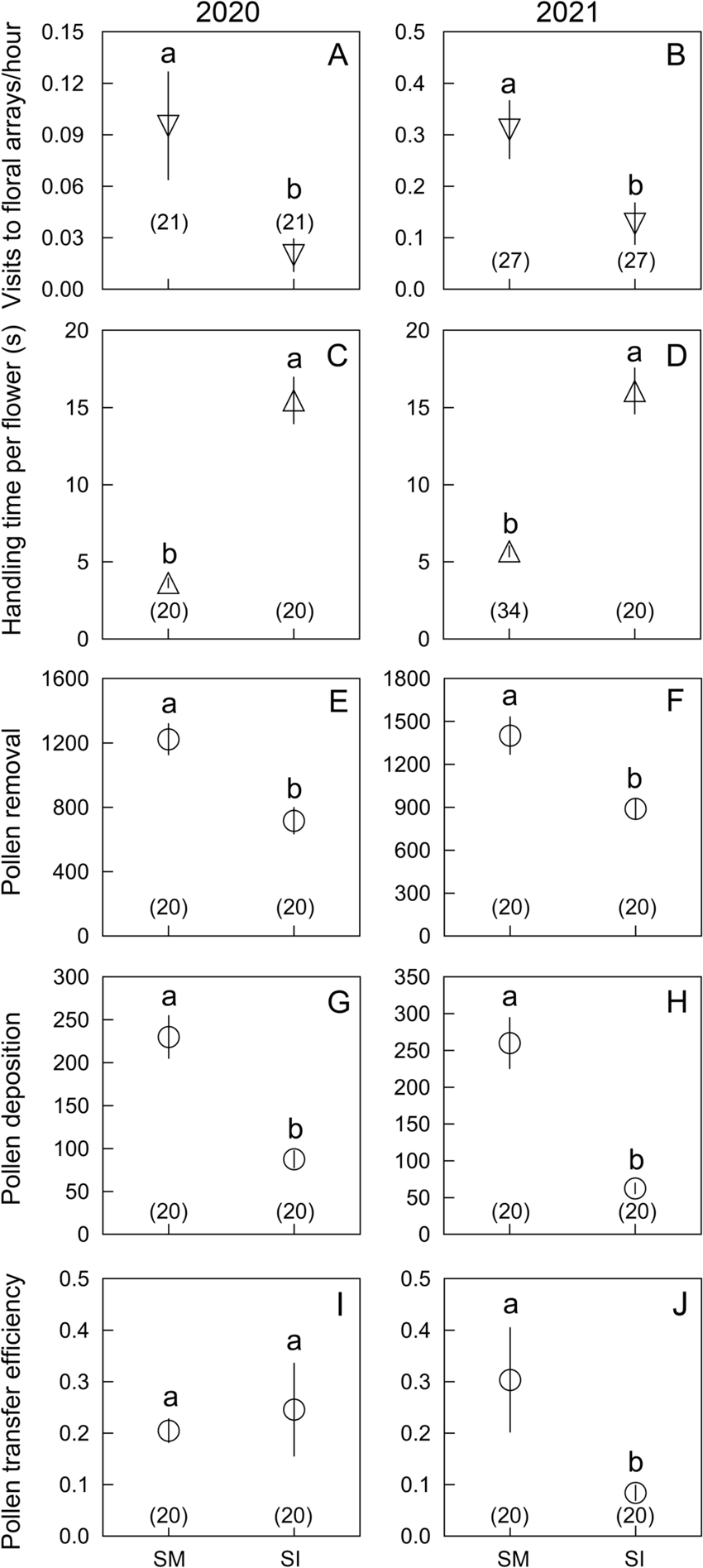
Foraging behaviour of *Apis cerana* in *Berberis julianae* flowers with experimentally immobilized stamens and controls over two years. Mean and standard errors (error bars) are presented, with different lowercase letters indicating significant differences between control (SM) and stamen immobilized (SI) flowers.

**Figure S6.**
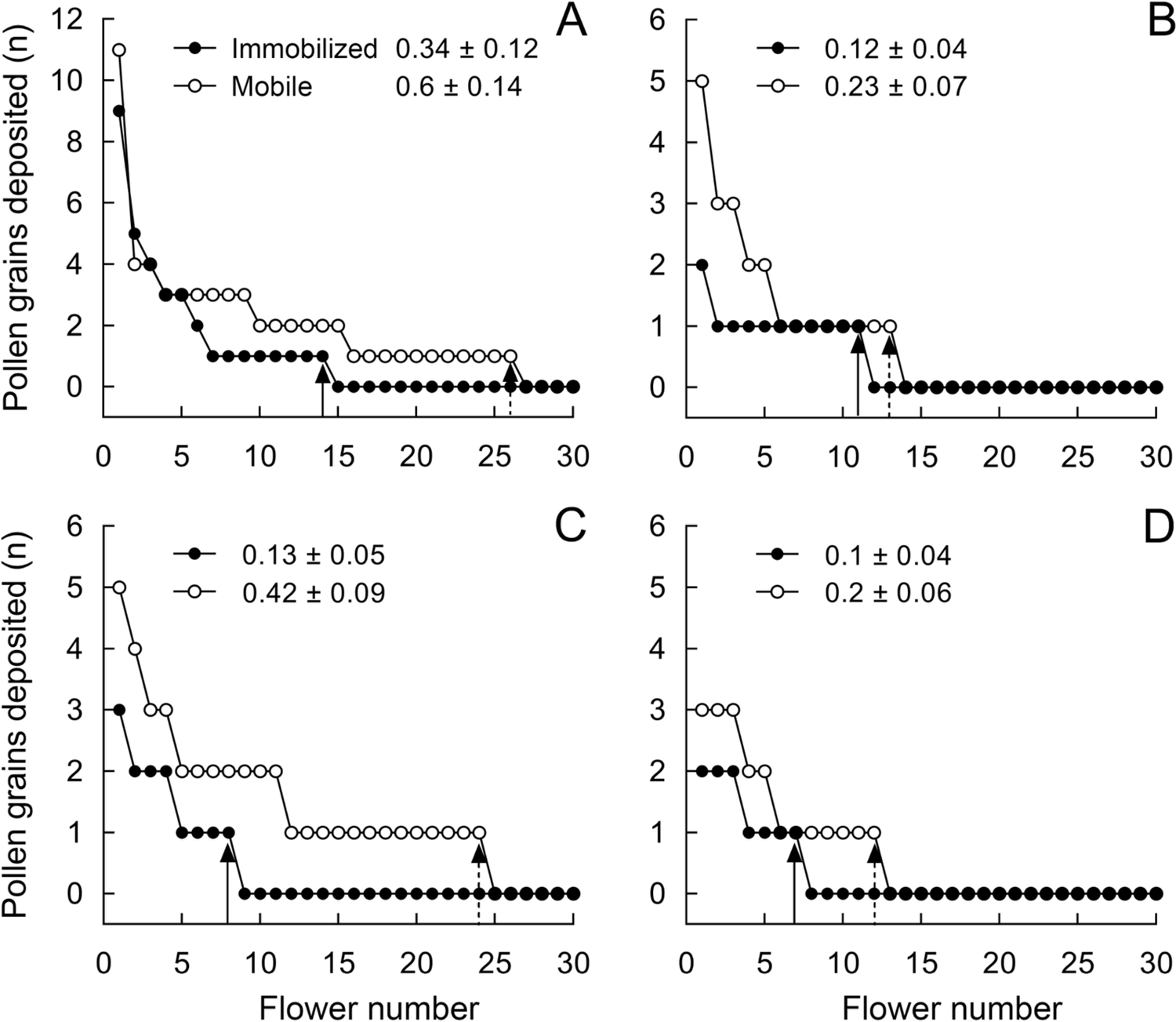
Number of stained pollen grains deposited on the stigmas of control flowers with mobile stamen (open circles) and on stigmas of flowers with experimentally immobilized stamens (closed circles) in *Berberis jamesiana*. Mean pollen number (n = 100 flowers) and standard errors are given in each of four trials (A, B, C and D). Note that the other 70 zeroes are not shown, as each pollen-tracking test involved 100 pollen-recipient flowers to examine the effect of stamen movements on pollen dispersal.

**Table S1.** Floral traits (mean ± SE) and duration of stamen movements in *Berberis jamesiana*, *B. julianae*, and *B. forrestii*, and *Mahonia bealei*.. Different superscript letters above means indicate significant differences; n = number of sampled flowers. Flowers are larger and take relatively longer to move inward or back in *Berberis julianae* than in the other two species.

| Floral traits | *Berberis julianae* | N | *Berberis jamesiana* | N | *Berberis forrestii* | N | *Mahonia bealei* | N | Wald $\chi^2$ | P |
| --- | --- | --- | --- | --- | --- | --- | --- | --- | --- | --- |
| Corolla diameter (mm) | 12.75<sup>a</sup> ± 0.40 | 20 | 8.37<sup>b</sup> ± 0.25 | 20 | 6.34<sup>c</sup> ± 0.14 | 30 | | | 324.66 | **0.000** |
| Petal length (mm) | 6.10<sup>a</sup> ± 0.07 | 20 | 4.78<sup>b</sup> ± 0.07 | 20 | 2.47<sup>c</sup> ± 0.07 | 30 | | | 1406.29 | **0.000** |
| Anther-stigma distance (mm) | 1.59<sup>a</sup> ± 0.08 | 20 | 0.96<sup>b</sup> ± 0.07 | 20 | 0.94<sup>b</sup> ± 0.06 | 30 | | | 52.85 | **0.000** |
| Pistil height (mm) | 5.48<sup>a</sup> ± 0.06 | 20 | 3.34<sup>b</sup> ± 0.08 | 20 | 2.28<sup>c</sup> ± 0.06 | 30 | | | 642.982 | **0.000** |
| Stamen length (mm) | 4.81<sup>a</sup> ± 0.07 | 20 | 2.69<sup>b</sup> ± 0.09 | 20 | 2.67<sup>b</sup> ± 0.07 | 30 | | | 303.741 | **0.000** |
| Filament length (mm) | 3.33 ± 0.06 | 20 | — | | — | | | | | |
| Nectar volume per flower (μL) | 1.20 ± 0.03 | 20 | — | | — | | | | | |
| Pollen number per flower | 7323<sup>a</sup> ± 78 | 20 | 6810<sup>b</sup> ± 156 | 20 | — | | | | 8.478 | **0.004** |
| Ovule number per flower | 3.10<sup>a</sup> ± 0.07 | 20 | 2.0<sup>b</sup> ± 0.0 | 20 | 2.03<sup>b</sup> ± 0.03 | 30 | | | 495.75 | **0.000** |
| Inward movement time (s) | 0.44<sup>a</sup> ± 0.02 | 10 | 0.17<sup>b</sup> ± 0.02 | 7 | 0.23<sup>b</sup> ± 0.04 | 5 | 0.09<sup>c</sup> ± 0.01 | 10 | 225.22 | **0.000** |
| Interval time (s) | 62.21<sup>a</sup> ± 3.08 | 10 | 13.19<sup>c</sup> ± 0.55 | 7 | 21.48<sup>b</sup> ± 3.56 | 5 | 3.46<sup>d</sup> ± 0.71 | 10 | 466.46 | **0.000** |
| Outward movement time (s) | 227.70<sup>a</sup> ± 10.06 | 10 | 110.37<sup>c</sup> ± 6.64 | 7 | 155.31<sup>b</sup> ± 14.07 | 5 | 7.74<sup>d</sup> ± 1.96 | 10 | 456.48 | **0.000** |

**Table S2.**
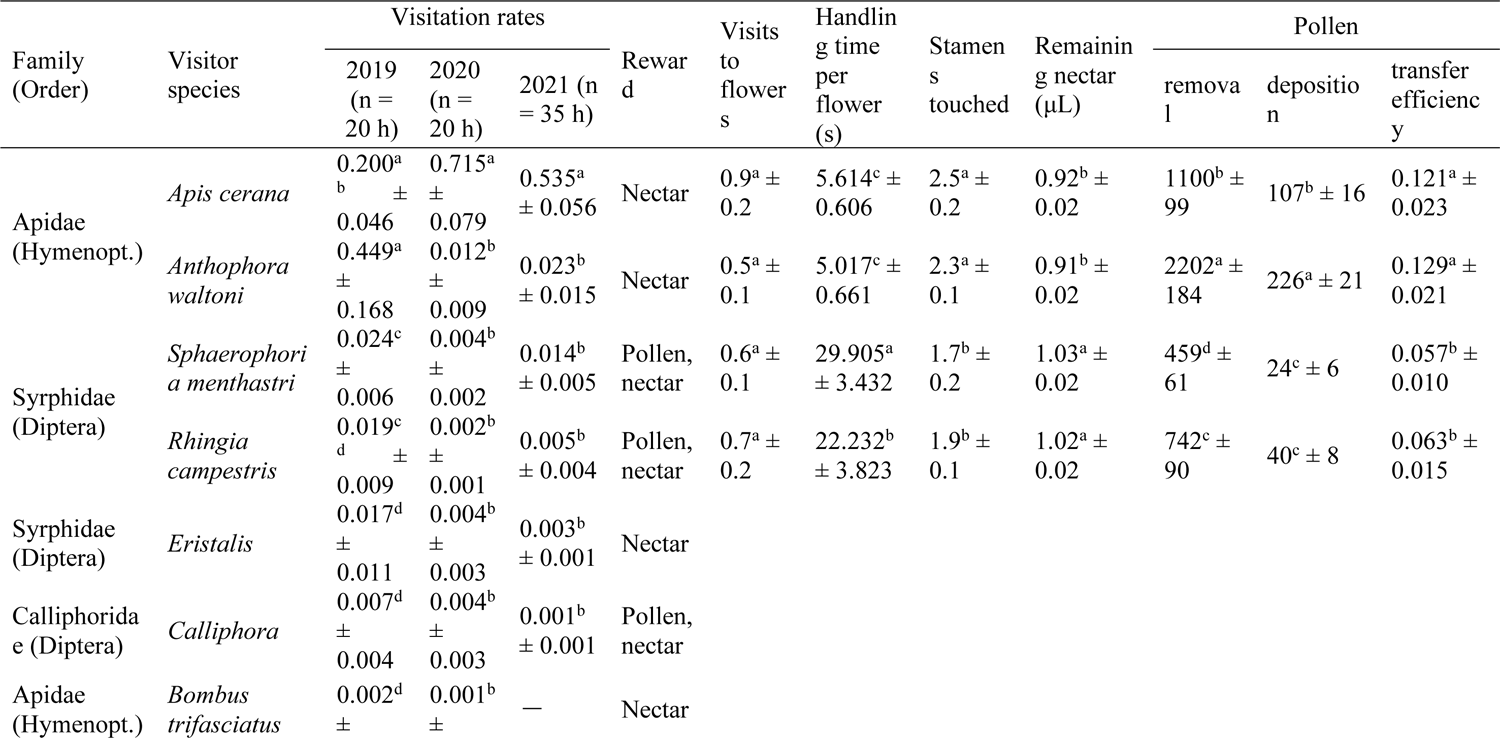
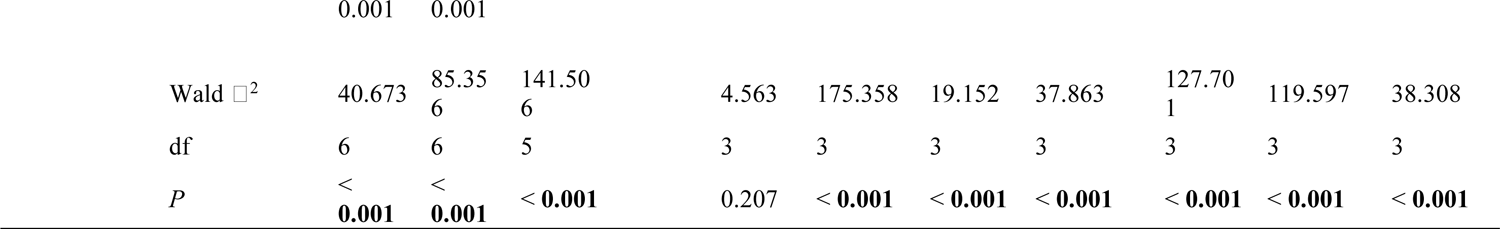
Behaviour and pollination roles of two species of bees and two species of flies visiting *Berberis julianae*. Visitation rates (visits per flower per hour, mean ± SE) in the field over three years, and visits per flower, handling time, number of stamens touched and nectar volume remaining per flower, pollen grains removed, pollen grains deposited and pollen transfer efficiency are compared among four insect species. Significant differences are shown with superscript letters.

**Table S3.**
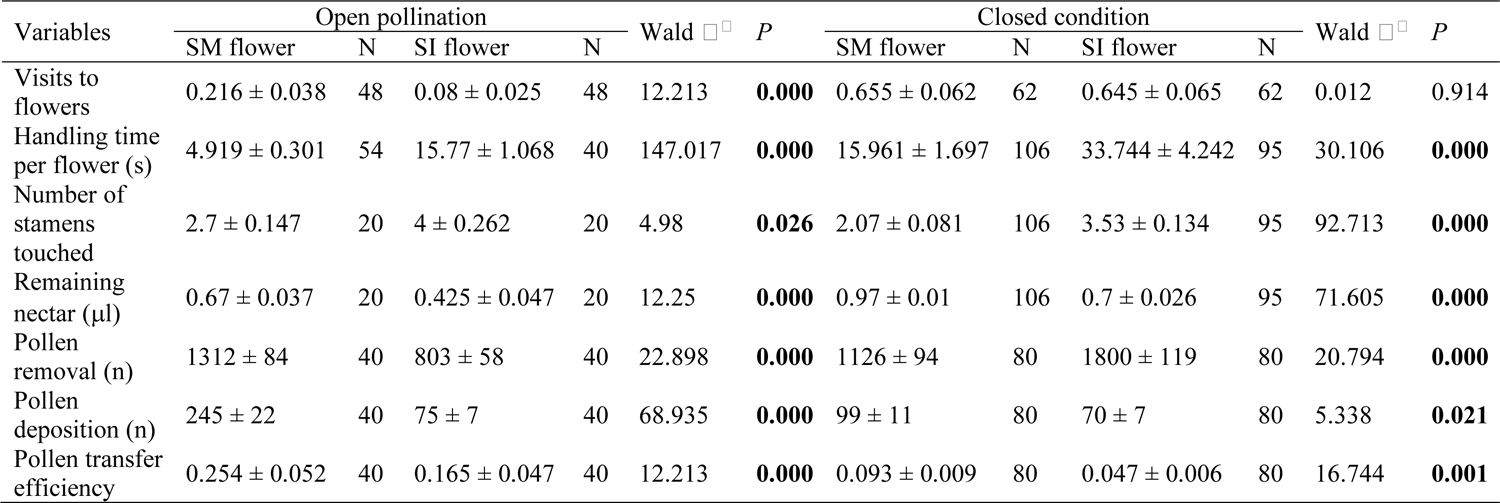
Foraging behaviour of visitors to *Berberis julianae* flowers with experimentally immobilized stamens and controls. For each measurement 20 flowers were sampled for each of four insect species under enclosed conditions (Fig. S1), but 40 flowers for the honeybee (*Apis cerana*) in open pollination conditions using data of two years. Mean values and standard errors of seven measured parameters are presented. Each comparison under enclosed conditions involves SM and SI flowers (N = number of flowers) visited by four visitor species including two bees (*A. cerana* and *Anthophora waltoni*) and two flies (*Rhingia campestris* and *Sphaerophoria menthastri*).

**Table S4.** Goodness of fit-tests confirming that *Berberis julianae* flowers with mobile stamens donated pollen to more recipient flowers (at >50 cm away) than did flowers with experimentally immobilized stamens.

| Pollen donors | Pollen-recipients/total flowers |  |  |  |
| --- | --- | --- | --- | --- |
|  | Distance < 50 cm | 50 cm < Distance < 100 cm | Distance > 100 cm | All |
| SM flowers | 25/338 | 19/434 | 6/222 | 50/994 |
| SI flowers | 10/395 | 4/338 | 0 | 14/733 |
| G | 21.555 | 22.435 |  | 35.805 |
| *P* | < **0.001** | < **0.001** |  | < **0.001** |

**Table S5.** Goodness of fit-tests confirming that *Berberis jamesiana* flowers with mobile stamens donated pollen to more recipient flowers (at >25 cm away) than did flowers with experimentally immobilized stamens. Flowers at short (0 - 25 cm) and greater distances (25 - 50 cm) from the pollen source receiving stained pollen and (/) total of sampled flowers in *Berberis jamesiana*.

| Pollen donors | Pollen-recipients/total flowers |  |
| --- | --- | --- |
|  | Distance < 25 cm | Distance > 25 cm |
| SM flowers | 13/60 | 7/60 |
| SI flowers | 9/46 | 1/56 |
| G | 0.164 | 15.037 |
| *P* | 0.685 | **0.0001** |

**Table S6.** Concentrations of berberine (mean ± SE) in different tissues of *Berberis julianae* under Generalized Linear Models. Different superscript letters indicate significant differences in that tissue (Wald χ2 = 46885.3, *P* < 0.001, df = 3).

| Tissues | Concentrations of berberine (mg/g) | N |
| --- | --- | --- |
| Leaf | 0.0519<sup>a</sup> ± 0.0001 | 10 |
| Petal | 0.0357<sup>b</sup> ± 0.0001 | 10 |
| Pollen | 0.0202<sup>c</sup> ± 0.0005 | 7 |
| Nectar | 0.0 ± 0.0 | 10 |

**Video 1.** Showing that stamens of *Berberis julianae* become touch-insensitive after the flower pedicels had been immersed in 75% alcohol for 35 minutes.

**Video 2.** Showing that stamens of *Mahonia bealei* become touch-insensitive after the flower pedicels had been immersed in 75% alcohol for 30 minutes.

**Video 3.** Under enclosed conditions (Fig. S1D), individual syrphid flies (*Sphaerophoria menthastri*) took up nectar from a *Berberis julianae* flower with immobilized stamens for much longer, giving them time to touch four stamens, while leaving more quickly and touching only two stamens in a control flower with mobile stamens.

